# Shelf-Life Specific Moisture Variation in Chitosan of Genus *Fenneropenaeus* Distributed Along Arabian Sea, India

**DOI:** 10.1101/2022.05.15.491996

**Authors:** Sreelekshmi R S, Lincy Alex, Jean J Jose

## Abstract

Chitosan is a characteristic polysaccharide, naturally extracted from the crustacean’s shells. The stability and shelf-life of chitosan are affected by a few parameters, basically the moisture content. In this study, three species of shrimps such as Indian white shrimp (*Fenneropenaeus indicus*), banana shrimp (*F. merguensis*), and tiger prawn (*F. monodon*) were utilized for the extraction of chitosan. The extraction strategy included the method of demineralization, deproteinization, and deacetylation. Later the yield and moisture of chitosan were examined from three diverse species. The yield of shell waste ranged between 46% to 50% (on moist weight premise) and the chitosan was between 18.0 to 20.2%. Moisture content always plays a critical part in chitosan shelf life and stability and was between 5.2% to 6.8 %.

## INTRODUCTION

Seafood, which can be found in a wide variety of products, is a delicacy in many coastal zones. Harvested products are packaged and normally processed by the seafood industry. In the course of processing, meat is only taken when the head and shell of molluscs and crustaceans are generated as waste. The consequence is the production of a large quantity of shell waste on a global scale. Shellfish waste constitutes many products of commercial value, such as chitin, chitosan, calcium carbonate, carotenoids, and proteins. The shrimp biowaste in the tropical region contains 10-20 % calcium, 30-65% protein content, and 8-10% chitin on a dry basis (Toan, 2009). Processing shellfish waste is an important source of wealth (Suryawanshi et al., 2019). A cost-effective solution to this problem is the recycling and reusing of shell wastes and the extraction of commercially viable substances like chitin. Chitin on its own has various applications. This can further be deacetylated to form chitosan which has a wide range of uses (Kumar, 2000).

Chitin (-(1–4)-poly-N-acetyl-D-glucosamine) is the most prevalent amino polysaccharide polymer discovered in nature, second only to cellulose. This material is found in the exoskeletons of insects, crustaceans, and fungal cell walls (Ma et al., 2014). Chitin is the principal structural component of clam, crab, and shrimp exoskeletons, as well as the cell walls of fungus. It is found in nature as organised macro fibrils. For biological reasons, chitin is routinely converted to its deacetylated derivative, chitosan (Divya et al., 2014). Chitosan is created by removing chitin of enough acetyl groups (CH3-CO), resulting in a linear chain of acetylglucosamine groups.

Chitosan (figure 1) is a biopolymer that degrades naturally and is converted to basic, non-toxic components by enzymes. In vivo, lysozyme, a non-specific protease present in all mammalian tissues, degrades chitosan mostly by producing non-toxic oligosaccharides that can be ejected or incorporated into glycosaminoglycans and glycoproteins. Invitro degradation of chitosan through oxidation, chemical or enzymatic hydrolysis reactions are commonly used methods for the preparation of low molecular (Ma et al., 2014). Chitosan exhibits numerous physicochemical properties such as naturally renewable sources, non-allergenic, antimicrobial, biocompatible, non-toxic, and biodegradable. Due to these integrated interesting properties, this biopolymer attracts great interest in a broad range of research fields, including the use in biomedicine, food, textile, and various chemical industries (Divya et al., 2014). In the food business, chitosan is employed as a binder, gelling, thickening, and stabilising agent in the chemical treatment of waste water.(Kumari & Rath, 2014). Despite its unique properties, chitosan is proving its efficacy in a variety of dosage forms, including bio adhesive nature, hydrophilic macromolecule drug carrier, effective carrier in drug targeting to the brain, transdermal films, and wound healing biodegradable grafts, hyperlipidaemic, an antimicrobial and stabilising constituent of liposomes, and wound healing biodegradable grafts. Chitosan has the potential to be a viable pharmaceutical excipient(Puvvada et al., 2012).

**Figure 1.**
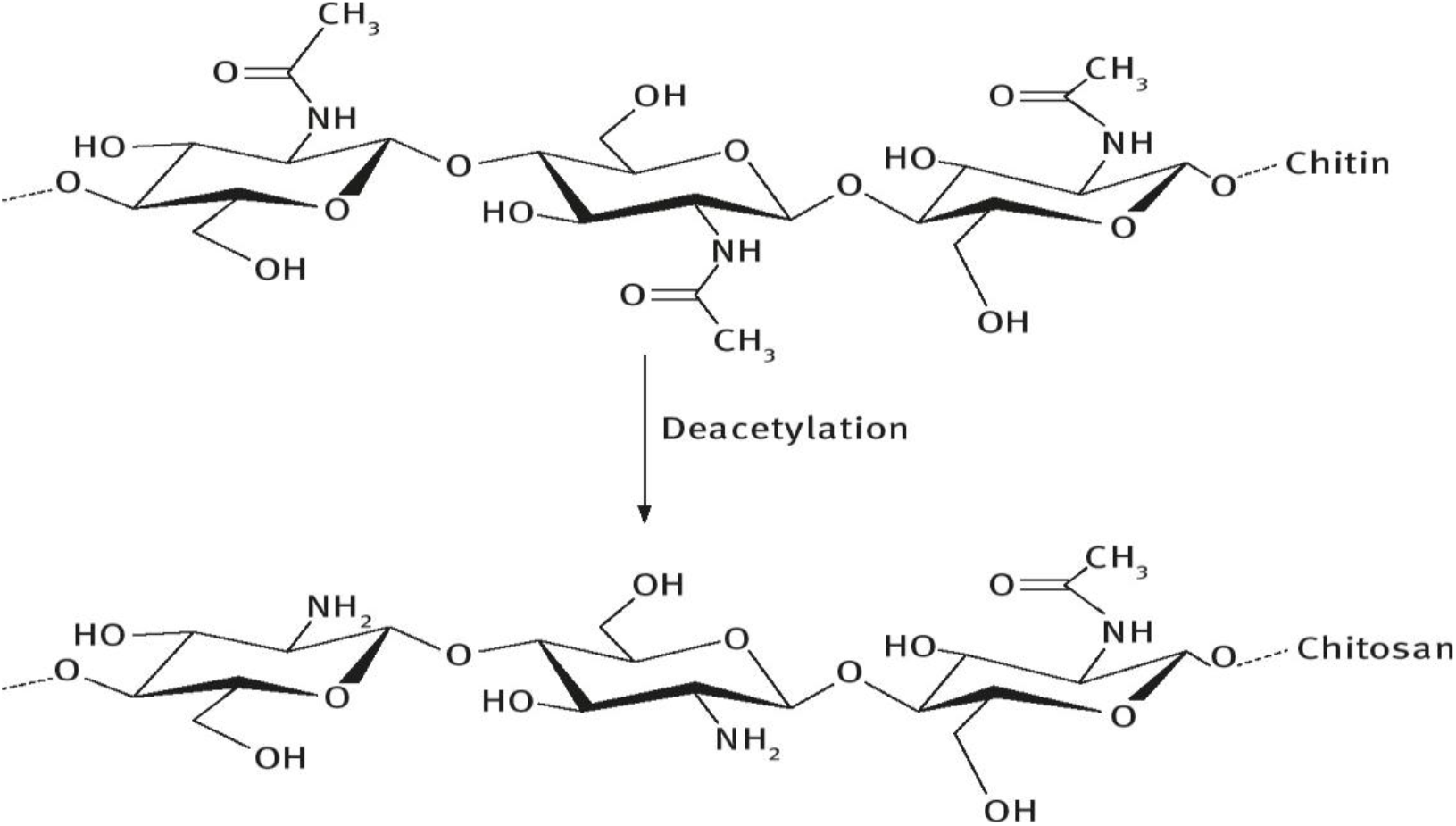
Chitin undergo deacetylation to form chitosan. (Nilsen-Nygaard et al., 2015)

Because of their exceptional biocompatibility, biodegradability, non-toxicity, chelating, and adsorption power, chitin and its derivative chitosan are of economic importance. (Casadidio et al., 2019). Chitosan has a wide range of uses in biotechnology, the food and pharmaceutical industries, cosmetics, environmental engineering, agriculture, and aquaculture (Vani & Stanley, 2013).

Chitosan is quite sensitive to environmental factors like moisture and temperature, therefore it’s best to keep it at room temperature and relative humidity. The relative humidity of the surrounding environment has a significant impact on the presence and distribution of moisture in the chitosan substance (Patria, 2013). Temperature, in addition to relative humidity, has an impact on moisture content in chitosan-based systems. The loss of moisture (dehydration of chitosan powder) was found to be substantial when exposed to high temperatures (40 °C)) (Viljoen et al., 2014).

The study aimed to compare the variations of moisture content related to shelf life of chitosan extracted from three species of *Fenneropenaus* and commercial type chitosan for its wide utilization.

## MATERIALS AND METHOD

Fresh local Indian white shrimp (*Fenneropenaeus indicus*), banana shrimp (*Fenneropenaeus merguensis*), and tiger prawn (*Fenneropenaeus monodon*) were collected from local fish market of Kollam District, Kerala. Head and flesh were peeled and shells were obtained and sun dried after cleaning. The dried shells were packed and sealed in a zip-lock bag and kept at ambient temperature (27 ±2°C) for the experiment purpose.

### Reagents used

1. Sodium Hydroxide (0.5% w/v): 0.5 gm NaOH per 100 ml distilled water, 2. Hydrochloric acid (1.25N to 1.5N): for 100ml stock solution, 10.8ml HCl is measured and the volume is made up to 100 ml with distilled water. 3. 1% Acetic Acid: 1ml acetic acid per 100 ml distilled water. 4. Sodium Hydroxide (3% w/v): 3gm NaOH per 100 ml distilled water. 5. Sodium Hydroxide (42% w/v): 42gm NaOH per 100 ml distilled water (Tarafdar & Biswas, 2013) (Figure 2).

**Figure 2.**
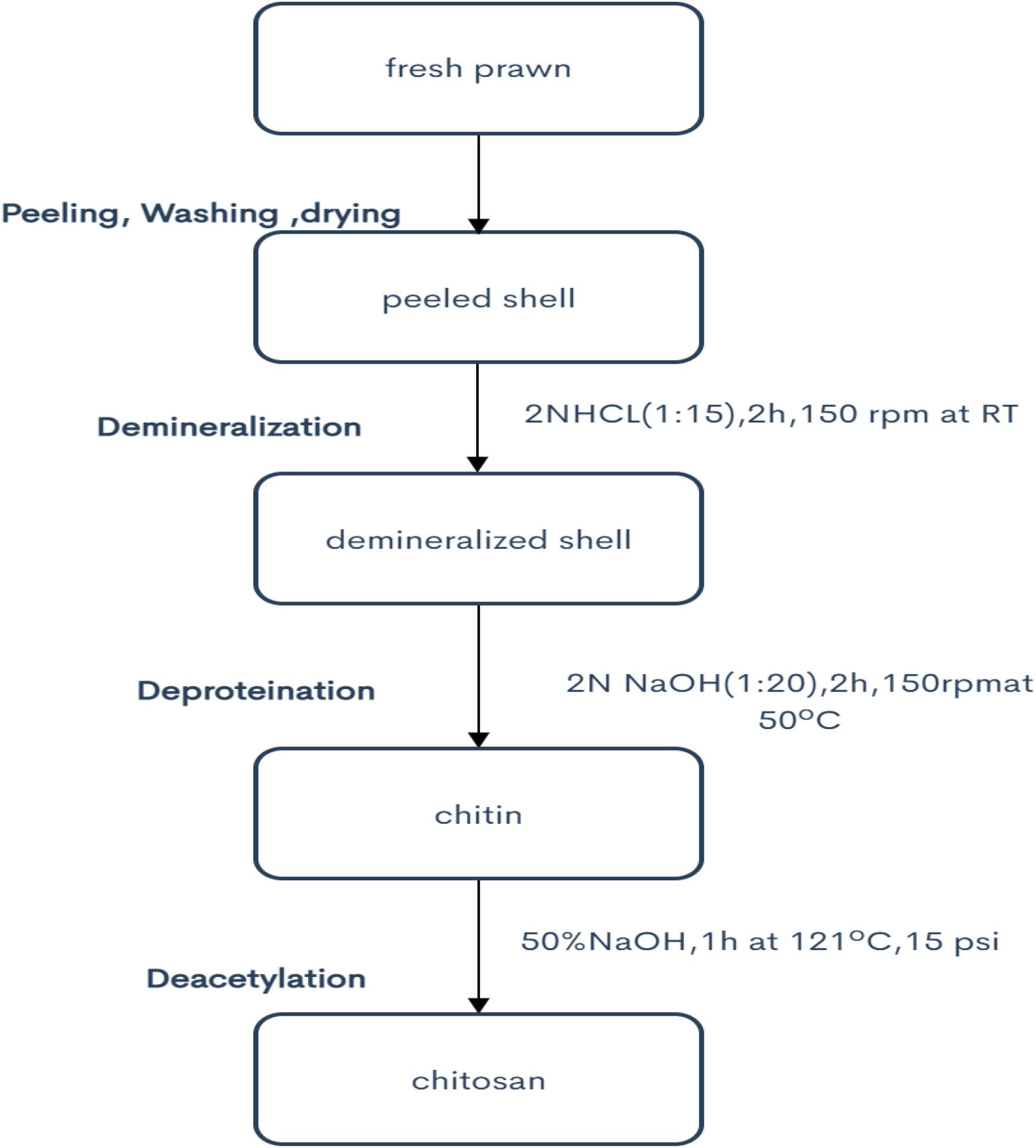
Extraction of chitosan from shrimp shell waste Based on (Tarafdar & Biswas, 2013) chemical extraction process.

### Preparation of chitin and chitosan

Chitin is made by a three-step chemical process that includes demineralization, deproteinization, and deacetylation, which leads to the development of chitosan. Figure 2 shows a schematic illustration of the extraction procedure.

### Demineralization

Shell wastes were demineralized in 1.5N HCl for 3 hours at room temperature (27±2 °C). The leftover HCl was rinsed away many times with distilled water until the pH was neutral.

### Deproteinization

After demineralization, the shells were deproteinized with 2N NaOH at 100°C for 30 minutes (Ushakumari & Ramanujan, 2012). The residual NaOH was rinsed away with distilled water many times to achieve the neutral pH. The filtered chitin was dried and finely powdered to enable for the deacetylation procedure. This approach helps to weaken the tertiary structure of proteins. For the elimination of any residual protein, the deproteinization procedure was repeated, 3 % NaOH was added to the sample at 100°C for 30 minutes.

The spent acid was removed and washed thoroughly with distilled water, with the pH being checked until it was near to neutral.

### Deacetylation

The acetyl groups in chitin from shell wastes were removed by treating it with 50 % aqueous NaOH at 95°C for 1.5 hours. After deacetylation, the alkali was removed and washed thoroughly with distilled water until the pH was less than 7.5, then dried at room temperature (31±2 °C).

### Estimation of chitosan yield

The weight of chitosan produced is measured and yield is calculated.

### Moisture content

Moisture content was analysed using hot air oven as per AOAC method (2000).The samples were dried for 3 hours at 105°C

Amount of moisture present in the chitosan plays a very important role in its stability and shelf life.

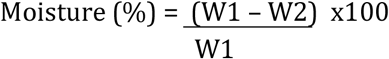

Where,

W1 = weight (g) of sample before drying
W2 = weight (g) of sample after drying, (*ANALYTICAL METHODS*, n.d.)AOA

## RESULTS AND DISCUSSION

### Yield of chitosan

Table I depict the yield of shell waste and chitosan obtained from different species of prawn. Chitosan yield ranges from 18-21 % and shell waste yield ranges from 46-50% (on wet weight basis). Chitosan resulted are illustrated in Figure 3.

**Table 1.**
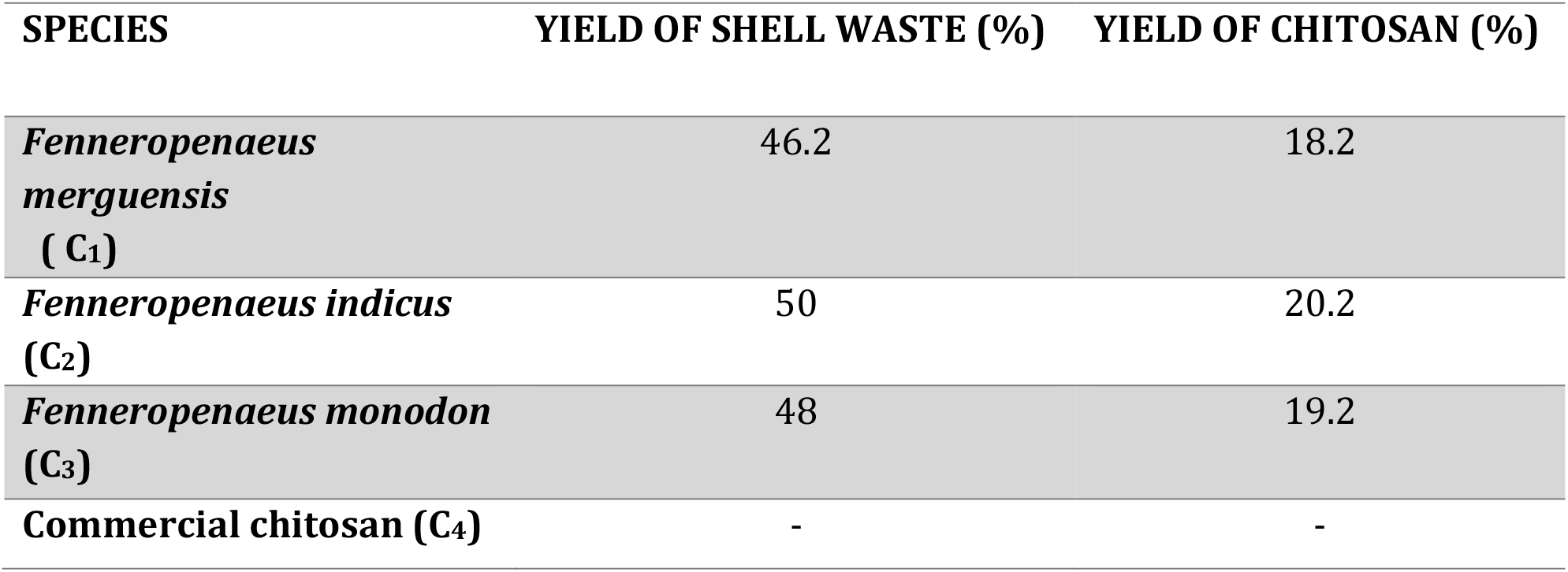
shell waste and chitosan yield from different species of prawn.

**Table 2 :** Moisture analysis of shell waste and chitosan.

| species | Shell waste moisture (%) | Moisture content in chitosan |
| --- | --- | --- |
| *Fenneropenaeus merguensis* (C<sub>1</sub>) | 67.2 | 5.2 |
| *Fenneropenaeus indicus* (C<sub>2</sub>) | 71.2 | 8.6 |
| *Fenneropenaeus monodon* (C<sub>3</sub>) | 69.3 | 6.7 |
| Commercial chitosan | - | 12.7 |

**Figure 3.**
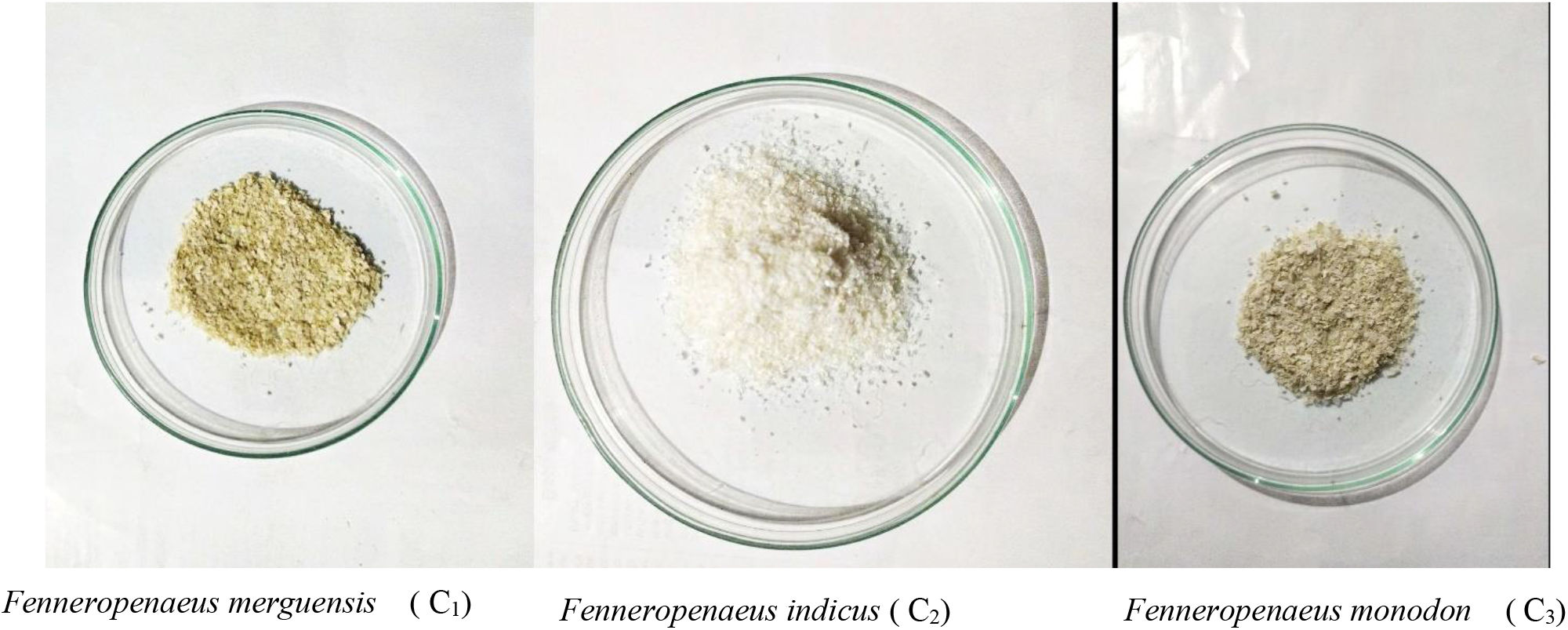
chitosan produced from different species of shrimp.

Results inferred the exoskeleton content yield of the shrimp species studied ranged from 45 to 55 %. It can be discussed with the earlier works of Lertsutthiwong et al., 2002 and Parthiban et al. which describes the shrimp shell waste had a maximum yield of 45.580%. The current study found that the shell waste production was between 46 to 50 percent, which is sufficient for further research. The yield of chitosan was reported to be around 20% by Brzeski (1982), 18.6 % by Alimuniar and Zainuddin (1992), and roughly 23 percent by No and Meyers (1989). Chitosan yields from three species ranged from 18 to 21% in this investigation, which is similar to Alimuniar and Zainuddin (1992) and No - Meyers results. The yield difference is related to response time, which has a favourable impact on yield. In general, yield values of chitosan decreases with increasing of heating temperature, probably due to the high temperature will cause the molecular chains of chitosan depolymerization process that eventually then cause decreases in molecular weight of chitosan (Patria, 2013).

### Moisture content

Table 1 shows the results of the moisture content of shell waste and chitosan. The moisture content of shrimp shell debris was found to be 71.6 % and 69.3 %, respectively, by Usha Kumari UN and Hossain MS. However, the moisture content of the shrimp shell wastes was found to be 67.2 %, 71.2 %, and 69.3 % in this investigation, which is similar to the previously mentioned values.

According to (Li et al., 1992) commercial chitosan products have less than 10% moisture, however (Rege et al., 1999) discovered that chitosan powder moisture levels ranged from 5% to 11% (w/w).

The moisture percentage of the chitosan samples ranged between 5.2 % to 12.7 %, indicating a modest variance in moisture content in chitosan pooled from each species. Table 3 showed earlier studies on the moisture content of shrimp shell debris with the current study. The amount of water absorbed is determined by the original moisture content as well as storage conditions, particularly temperature and relative humidity(Hossain & Iqbal, 2014).

**Table 3 :** Comparative analysis of moisture content of chitosan of previous and present studies.

| Species | Moisture content | Reference |
| --- | --- | --- |
| *Fenneropenaeus merguensis* (C<sub>1</sub>) | 5.2 | Present study |
| *Fenneropenaeus indicus* (C<sub>2</sub>) | 8.6 | Present study |
| *Fenneropenaeus monodon* (C<sub>3</sub>) | 6.7 | Present study |
| Commercial chitosan | 12.7 | Present study |
| *Fenneropenaeus indicus* | 7.67% | (Ushakumari & Ramanujan, 2012) |
| *Penaeus monodon* | 5% | (Divya et al., 2014) |
| *Aristeus alcocki* | 9.23% | (Ushakumari & Ramanujan, 2012) |
| **Fresh shrimp was procured from Mymensingh's local market.** | 7.69% to 8.25% | (Hossain & Iqbal, 2014) |
| **Commercial chitosan** | 7% to 11% | (Rege et al., 1999) |

The presence of absorbed water has a substantial influence on the flow properties and compressibility of solid chitosan-based formulations.(No & Prinyawiwatkul, 2009). When the moisture content of the material varies during storage, the physicochemical and mechanical features of chitosan-based systems may vary. The discrepancies in moisture content in this study are due to the samples’ lack of homogeneity. According to research done by (Natarajan et al., 2017) the amount of absorbed water in chitosan powder increased during storage, resulting in a decrease in water binding ability. Due to the formation of hydrogen bonds between the particles, moisture content of up to 6% (w/w) has been reported to promote particle binding during compression. According to recent studies, the typical moisture content of chitosan is less than 10%. Moisture content has a crucial effect in stability, particularly for chitosan’s shelf life (Ushakumari & Ramanujan, 2012). In this comparative analysis, naturally extracted chitosan has a moisture content below 10%, in which C2 shows the high moisture content of 8.6% and C1 with low moisture content of 5.2. But in case of commercial chitosan, the result showed a higher rate of moisture content of 12.7%.

As the rate of moisture content increases the stability and shelf life of chitosan decreases. From the above result, it is understood that the commercial chitosan is easily vulnerable to degradation than naturally extracted ones. From the naturally extracted chitosan, chitosan from *Fenneropenaeus merguensis* (C_1_) is highly stable and has a long shelf life, which can be stored for a long time at ambient temperature.

## CONCLUSION

Shrimp shells were treated and extracted for chitosan using standard protocol. Mild variations in moisture concentration were recorded from species specific naturally extracted chitosan when compared with commercial chitosan which recorded considerably a high rate. Hence product specific naturally extracted chitosan is highly stable with long shelf life and can be highly recommended for biomedical applications, food industries and industries.

## AKNOWELDGEMENT

Authors convey their sincere thanks to CEPCI (The Cashew Export Promotion Council) Laboratory and Research Institute, Kollam and Central Laboratory for Instrumentation and Facilitation (CLIF) University of Kerala, Thiruvananthapuram

